# Somatic piRNAs and Transposons are Differentially Regulated During Skeletal Muscle Atrophy and Programmed Cell Death

**DOI:** 10.1101/2021.03.02.433533

**Authors:** Junko Tsuji, Travis Thomson, Christine Brown, Subhanita Ghosh, William E. Theurkauf, Zhiping Weng, Lawrence M. Schwartz

## Abstract

PiWi-interacting RNAs (piRNAs) are small single-stranded RNAs that can repress transposon expression via epigenetic silencing and transcript degradation. They have been identified predominantly in the ovary and testis, where they serve essential roles in transposon silencing in order to protect the integrity of the genome in the germline. The potential expression of piRNAs in somatic cells has been controversial. In the present study we demonstrate the expression of piRNAs derived from both genic and transposon RNAs in the intersegmental muscles (ISMs) from the tobacco hawkmoth *Manduca sexta.* These piRNAs are abundantly expressed, are ~27 nt long, map antisense to transposons, are oxidation resistant, exhibit a uridine bias at their first nucleotide, and amplify via the canonical ping-pong pathway. An RNA-seq analysis demonstrated that 20 piRNA pathway genes are expressed in the ISMs and are developmentally regulated. The abundance of piRNAs does not change when the muscles initiate developmentally-regulated atrophy, but are repressed when cells become committed to undergo programmed cell death at the end of metamorphosis. This change in piRNA expression is associated with the targeted repression of several retrotransposons and the induction of specific DNA transposons. The developmental changes in the expression of piRNAs, piRNA pathway genes, and transposons are all regulated by 20-hydroxyecdysone, the steroid hormone that controls the timing of ISM death. Taken together, these data provide compelling evidence for the existence of piRNA in somatic tissues and suggest that they may play roles in developmental processes such as programmed cell death.

**Author Summary:** piRNAs are a class of small non-coding RNAs that suppress the expression of transposable elements, parasitic DNA that if reintegrated, can harm the integrity of the host genome. The expression of piRNAs and their associated regulatory proteins has been studied predominantly in germ cells and some stem cells. We have found that they are also expressed in skeletal muscles in the moth *Manduca sexta* that undergo developmentally-regulated atrophy and programmed cell death at the end of metamorphosis. The expression of transposons becomes deregulated when the muscles become committed to die, which may play a functional role in the demise of the cell by inducing genome damage. piRNA-mediated control of transposons may represent a novel mechanism that contributes to the regulated death of highly differentiated somatic cells.

## Introduction

Small silencing RNAs are powerful regulators of gene expression. They can lead to epigenetic silencing of transcription, transcript degradation, and inhibition of mRNA translation. The best characterized class of small silencing RNAs are the microRNAs (miRNAs)(1–3). miRNAs are ~22 nucleotides (nt) long, are ubiquitously expressed, and can repress their target transcripts through seed-based partial complementarity (4–6).

The most recently discovered group of small silencing RNAs are PIWI-interacting RNAs (piRNAs). piRNAs are 23-35 nt in length and predominantly expressed in the germline of animals including humans (7–10). They guide the PIWI clade of Argonaute proteins to silence transposons and other selfish elements, and protect the integrity of the germline genome (reviewed in Ozata, Gainetdinov et al., 2019)(11).

The biogenesis and function of piRNAs has been well studied in the fruit fly *Drosophila melanogaster* (12). piRNAs are processed from long transcripts that can be up to hundreds of kilobases long that are transcribed from discrete genomic loci called “piRNA clusters” (13–15). piRNAs are amplified via reciprocal target cleavages by PIWI proteins, a mechanism known as the ping-pong cycle (13, 16). Because PIWI proteins cleave the phosphodiester bond between the nucleotides in the target RNA that pair with the 10^th^ and the 11^th^ nucleotides of the guide piRNA, and 3’ cleavage products is subsequently made into another piRNA, there is an enrichment of piRNAs that perfectly reverse complement each other in their first 10 nucleotides, the hallmark of the ping-pong cycle. This process typically creates piRNAs with a uridine residue as the first nucleotide of the primary piRNA, and complementarity over the first 10nt of post-transcriptionally amplified piRNAs (13, 17–19).

piRNAs predominantly target transposons, retroviruses and other “selfish” genetic elements. In the absence of piRNAs, transposons can mobilize resulting in double-stranded DNA breaks in the germline genome leading to infertility (20, 21). Consequently, the expression and function of piRNAs has been most extensively studied in germ cells, gonadal somatic cells, and certain progenitor cells (13, 22, 23).

While there have been several reports demonstrating the presence of both piRNAs and the associated protein machinery in non-gonadal cells, their role in cellular processes has been a controversial subject (24–28). Somatic piRNAs have been observed broadly in arthropods where they may provide genome defense against transposable elements and viruses (29). In agreement with this hypothesis, piRNAs from fat body and midgut cells elicits antiviral response against nucleopolyhedrovirus in the silkmoth *Bombyx mori* (30). In addition, piRNAs derived from endogenous viral elements system in the mosquito *Aedes aegypti* help maintain long-lasting adaptive immunity (31, 32). Data has been generated suggesting that piRNAs may act in the nervous system and influence transposon activity, learning and memory. For example, in *Drosophila,* the piRNA pathway proteins Aub and Ago-3 regulate transposon expression and mobilization in the mushroom bodies, brain structures that regulate memory formation and cognitive function (33). These proteins are also found in specialized structures within glial cells in the adult *Drosophila* brain where they appear to repress transposon activity (34). In the sea slug Aplysia, a specific piRNA has been shown to modulate synaptic plasticity and memory (35, 36).

In the current study we provide substantial data demonstrating the presence of piRNAs and their synthetic machinery in the intersegmental muscles (ISMs) from the tobacco hawkmoth *Manduca sexta.* We further demonstrate that transposon expression becomes deregulated when these cells become committed to undergo programmed cell death at the end of metamorphosis.

The ISMs are a classical model system for skeletal muscle atrophy and death (37–39). These muscles are composed of giant syncytial cells, each of which is about 5 mm long and up to 1 mm in diameter. The ISMs are used by the larvae to crawl and by the developing adult moth to eclose (emerge) from the overlying pupal cuticle at the end of metamorphosis. Three days before eclosion (day 15 of the normal 18-day period of pupal-adult development), the ISMs initiate a program of atrophy that results in a ~40% loss of muscle mass prior to eclosion. This atrophy is non-pathological and the muscles retain normal physiological properties such as resting potential and force per cross-sectional area (40). Late on day 18, coincident with adult eclosion, the ISMs initiate programmed cell death (PCD) and die during the subsequent 30 hours. In fact, the term PCD was coined by Lockshin and Williams in 1965 to describe the death of these specific cells (41). PCD is a fundamentally different program than atrophy and results in the complete destruction of the contractile apparatus, loss of the resting potential, and enhanced autophagy (40, 42). The dying cells are not phagocytosed and instead activate the molecular machinery required for both cellular destruction and nutrient recycling (42, 43). The developmental timing of both atrophy and death is controlled by circadian declines in the circulating titer of the insect molting hormone 20-hydroxyecdysone (20E) (44). Judiciously timed administration of exogenous 20E can prevent atrophy or death, but once either of these programs has been initiated, it cannot be altered or delayed by 20E treatment.

Several studies have demonstrated that ISM PCD requires *de novo* gene expression, and numerous death-associated genes have been identified (42, 45, 46). During the transition from atrophy to death, some ISM transcripts display significant changes in stability and translatability that can be localized to their 3’ untranslated regions (UTRs) (47). Indeed, direct testing has demonstrated that the specific microRNAs can regulate the translation of specific death-associated transcripts (42).

To further examine the potential role(s) of small silencing RNAs in the control of ISM atrophy and death, we performed RNA-seq with the small RNAs isolated from the ISMs each day of development from before the initiation of atrophy (day 13) until when the muscles were committed to die (day 18) (42). We also analyzed ISMs from animals that had been injected with 20E on day 17, a treatment that delays cell death on day 18. In addition to miRNAs, we found that the ISMs also contain high levels of piRNAs that are: ~27 nt long, map antisense to transposons, are oxidation resistant, exhibit a uridine bias at their first nucleotide, and amplify via the canonical ping-pong pathway. In addition, the ISMs express the genes required for piRNA synthesis and activity. When the ISMs become committed to die, there is a both a loss of piRNAs and the concurrent deregulation of transposable elements, with repression of some retrotransposons and the induction of DNA transposons. The expression of piRNAs and transposable elements are under the control of 20E. Thus, piRNAs are expressed in somatic tissues where they may regulate developmental processes such as PCD.

## Results

### piRNAs in the intersegmental muscles prior to atrophy

On day 13 of the pupal-adult development the ISMs are fully functional and have yet to initiated either the atrophy or PCD (44). We isolated the 18-30 nt small RNAs at this time point, and either cloned and sequenced them directly, or first oxidized the sample to render small RNAs that are not 2’-O-methylated at their 3’ termini (e.g., miRNAs) unclonable, thus enriching for piRNAs (48). Detailed mapping results for all small RNA-seq libraries are provided in Table S1.

We obtained 17.8 million and 11.4 million reads in the control and oxidized day 13 small RNA libraries respectively, of which 71.7% and 75.8% mapped to the *Manduca* genome. When the mapped reads in the unoxidized small RNA library were partitioned by size, we observed a bimodal distribution, with peaks at 22 and 27 nt (Figure 1). Among the reads shorter than or equal to 23 nt, we annotated 77.7% (78.1% for total reads) as miRNAs (42). In sharp contrast, among the reads longer than 23 nt, only 0.4% were miRNAs and instead, these RNAs displayed a strong 5’ uridine bias and a weak adenine signal at the 10th position (Figure S1).

**Figure 1:**
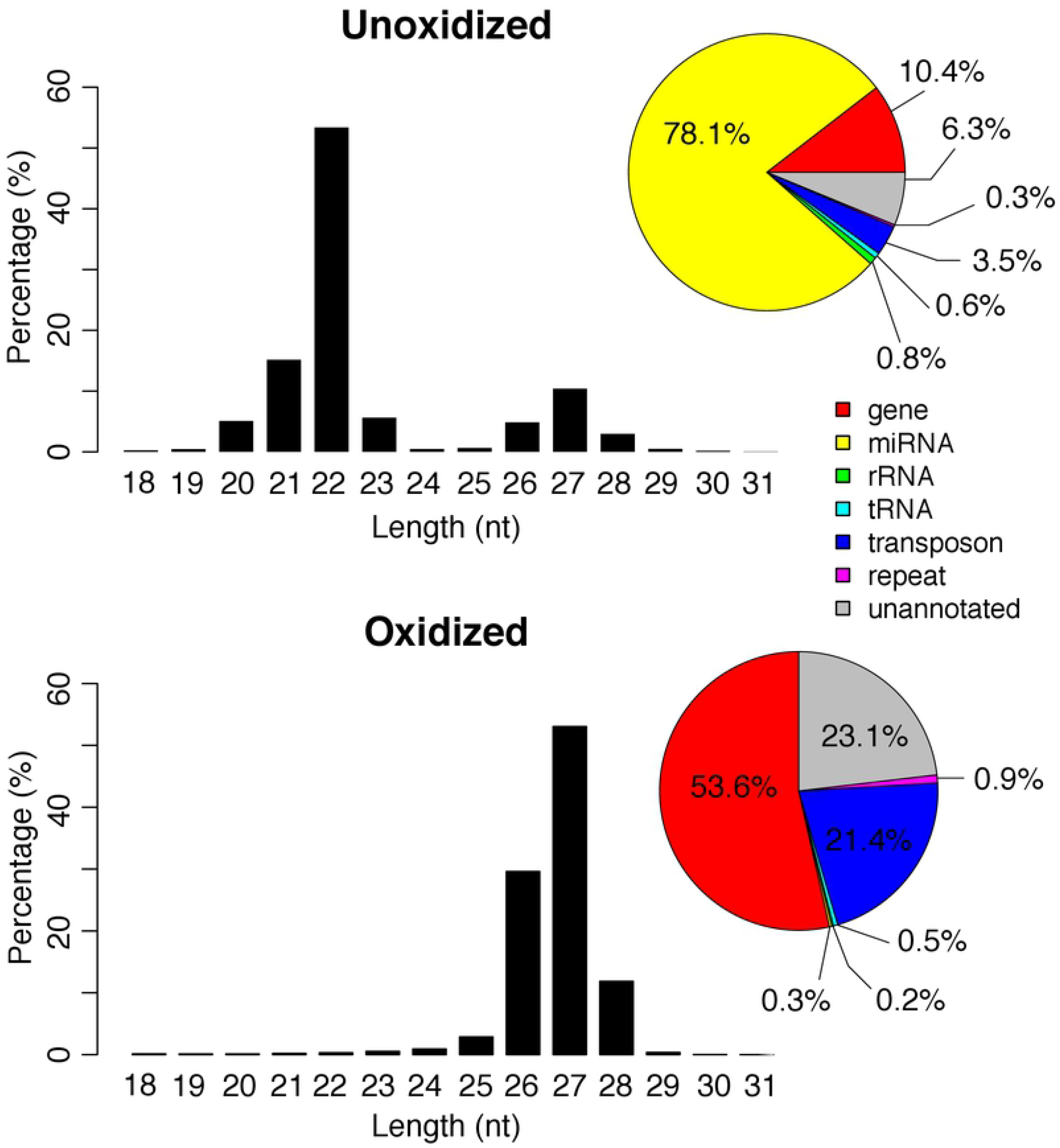
Length and mapping locations of *Manduca* piRNAs. Length distribution of the small RNAs in day 13 unoxidized (top) and oxidized (bottom) libraries. Genomic annotations of the locations where the reads mapped are summarized in pie charts (right).

Oxidation resulted in an almost complete loss of the 22 nt peak and 98.6% of the reads in the oxidized library were >23 nt (Figure 1). The majority of these reads started with a uridine base at the 5’ position (Figure S1). Only 0.3% of the reads in this library were identified as miRNAs. In contrast, 53.6% of the reads mapped to genes, 21.4% mapped to transposons, and 23.1% mapped to unannotated regions of the genome (Figure 1). These percentages were much higher than the reads ≤ 23 nt reads in the unoxidized library (0.8%, 0.3%, and 0.6% respectively), and comparable to the reads >23 nt in the unoxidized library (9.7%, 3.2%, and 5.6% respectively). Based on size and resistance to oxidation the 27nt peak appears to represent piRNAs.

### piRNA expression changes during the ISM development

In addition to the libraries on day 13, we generated and sequenced unoxidized small RNA libraries from seven more time points of ISM development: days 14, 15, 16, 17, 18, 1-hour post-eclosion (PE) and 20-hydroxyecdysone (20E) treated animals. Similar to what was seen with the day 13 unoxidized sample, the small RNAs from the other time points also displayed a bimodal distribution of small RNAs, with the majority of the ~22nt sequences mapping to miRNAs (Figure S1). In all of these samples, there was a strong bias for uracil in the first nucleotide, especially for the piRNAs (reads >23 nt).

piRNAs abundance from each stage was normalized by the sequencing depth and calculated as “parts per million” (ppm). The expression of both genic and transposable element piRNAs gradually declined from day 13 to day 16, increased sharply on day 17, and then declined dramatically on day 18 (Figure 2A). piRNA abundance was elevated in the one-hour post-eclosion (PE) sample. While treatment with 20E on day 17 delays ISM death (44) it did not significantly alter piRNA abundance in the muscles relative to the corresponding PE timepoint. Interestingly the number of piRNAs mapped to genes was more abundant than those mapping to transposons (Figure 2A).

**Figure 2:**
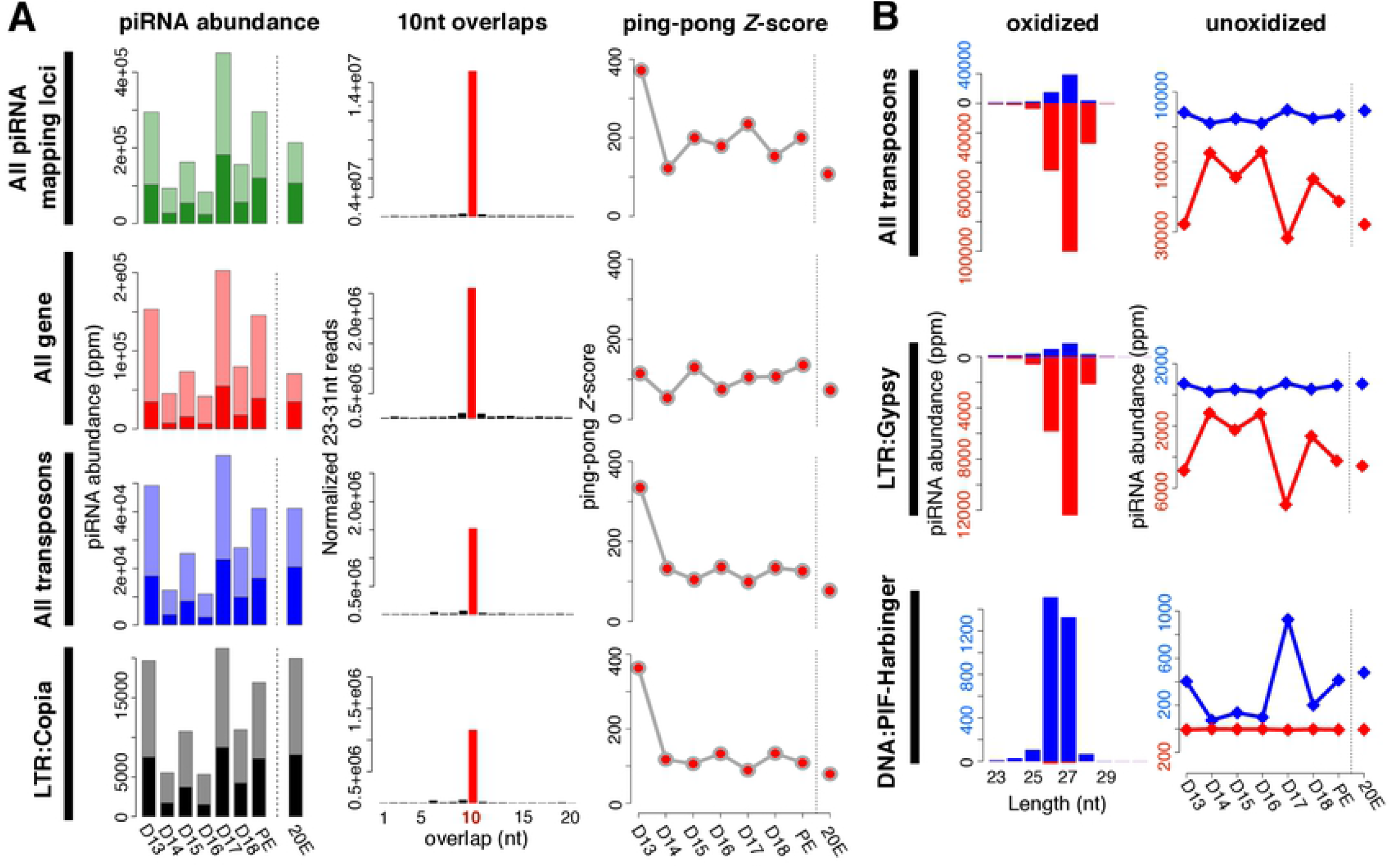
Characteristics of piRNAs expressed in the ISM. **A**: Changes in piRNA abundance during the ISM development (left); the frequency of 5’-5’ 10nt overlaps between piRNA pairs (middle); and the corresponding ping-pong Z-scores during development (right). The shaded bars on the left panels indicate the fraction of piRNAs that are amplified via the ping-pong amplification loop. Almost all of the Z-scores were statistically significant (red dots). **B**: Size distribution and strand bias of piRNAs mapped to transposons (left), and the change in piRNAs expression during the ISM development (right). Blue reflects sense strand bias while red is indicative of antisense. As examples, piRNAs mapped to DNA/PIF-Harbinger are in the upper panel while those mapping to LTR/Gypsy are on the bottom panel.

Of the 64 known transposon families in *Manduca,* 34 generated piRNAs that mapped to the genome. Among them, piRNAs for 27 transposon families mirrored the pattern of expression observed for all piRNAs and transposon-mapping piRNAs (e.g. LTR:Copia in Figure 2A) during ISM development. In contrast, the expressions of piRNAs mapped to five transposon families (two DNA transposon families: P and TcMar; and two LINE families: CR1 and I; and one SINE family: tRNA) were reduced gradually during the ISM development (Figure S2).

### piRNAs predominantly map antisense to transposons

The majority of piRNAs in flies that map to transposons are antisense to the mRNAs in order to guide the PIWI proteins Aub and Piwi to repress transposon expression (7). Consequently, mutations in piRNA pathway proteins that alter this antisense bias, such as Qin, lead to transposon de-repression (49). We examined the strand bias of transposon-mapping piRNAs at each stage of ISM development. Of the 34 transposon families with mapped piRNAs, we found that the majority of them (28 families) displayed strong antisense bias, while the remaining 6 families displayed a sense bias. The antisense piRNAs to transposons tend to be 27nt in length (e.g. LTR:Gypsy in Figure 2B), and the sense piRNAs to transposons tend to be either 26nt or 27nt (e.g. DNA:PIF-Harbinger in Figure 2B). Although the relative abundance of piRNAs was regulated during the course of ISM development their length distributions were constant.

### piRNAs expressed in the ISM amplify via the ping-pong amplification loop

Following transcription from piRNA clusters, the primary transcripts are processed and cyclically amplified by a mechanism known as the “ping-pong amplification loop” (7, 50). The secondary piRNAs generated via ping-pong tend to have a 10nt 5’ end overlap with other piRNAs on the opposite strand. This ping-pong activity is quantified by calculating the frequency of 5’-5’ overlaps between piRNA pairs, normalized as a Z-score (51).

We detected high Z-scores at all stages examined. For example, piRNAs from day 17 had an overall Z-score of 234.6 (Figure 2A). piRNAs from both transposons and genic sequences were amplified by ping-pong as demonstrated by Z-scores of 98.7 and 105.5 respectively. Z-scores were high on day 13, fell precipitously during the next three days, and then rose on day 17 and remained high for the rest of development, which agrees well with the abundance of piRNAs in the tissue. It should be noted that these developmental changes in Z-score were not artifacts arising from variations in sequencing depth since these same patterns were retained when we down sampled to 8 million reads per stage prior to our analysis.

As part of our analysis, we computed the Z-scores for piRNAs that mapped to each transposon class. Out of 34 transposon families with detectable piRNAs, 20 families displayed a high ping-pong signature throughout ISM development (e.g. LTR:Copia in Figure 2A), and 14 families displayed statistically significant Z-scores at least transiently. Intriguingly, in 12 out of 14 transposon families, the ping-pong signature peaked on day 17 in advance of the commitment of the ISMs to die. These data support the hypothesis that ISM piRNAs amplify via the traditional ping-pong amplification loop.

### Genic piRNAs preferentially map to 5’UTRs

Combining the RepeatMasker result with our own gene annotation, 8,633 (45.9%) of the 18,806 genes in *M. sexta* contain transposons within their introns. From the mapping results with our oxidized small RNA-seq library, we observed that piRNAs (24.38 ppm per intron in median) fell into the transposon-derived introns from 6,902 genes (80.0% of 8,633 genes containing transposons). piRNAs also mapped to introns that did not contain transposons, although the abundance was far lower (1,731 out of 18,806 genes (9.2%)). Compared to the abundance of piRNA reads on transposon-derived introns, there were very few non-transposon introns (2.29 × 10^−2^ ppm per intron in median).

It has been demonstrated that genic piRNAs tend to map to the sense orientation of 3’UTRs in germline and somatic cells (52–54). However, in the ISMs, we observed that the genic piRNAs tended to map to the antisense strand and preferentially within the 5’UTRs rather than the 3’ UTRs or coding sequences (CDSs). The strand bias patterns of the genic piRNAs uniquely mapped to 5’UTRs, CDSs, 3’UTRs, and introns were the same as those with all genic piRNA reads. Focusing on top 25% of the genes highly enriched with piRNAs, we next investigated the enrichment of piRNAs in 5’UTRs, CDSs, and 3’UTRs. In the oxidized small RNA-seq library, 5’UTRs tended to be more enriched with piRNAs mapped to the sense strands of genes as compared to other gene domains. In unoxidized small RNA-seq libraries, the overall trends were stably observed through all time points.

Considering the result that genic piRNAs tend to fall into the sense strands of 5’UTRs, we investigated the relative mapping positions of piRNAs in genes by examining the oxidized small RNA-seq library on day 13. Interestingly, when the piRNAs that mapped to highly expressed genes were plotted onto the gene map, there was a striking peak in the sense orientation in the 5’UTRs (Figure 3). (Only a modest number of piRNAs mapped to low abundance transcripts). In unoxidized small RNA-seq libraries, genic piRNAs tended to map to 5’UTRs and to form the peaks of mapped piRNAs (Figure S3).

**Figure 3:**
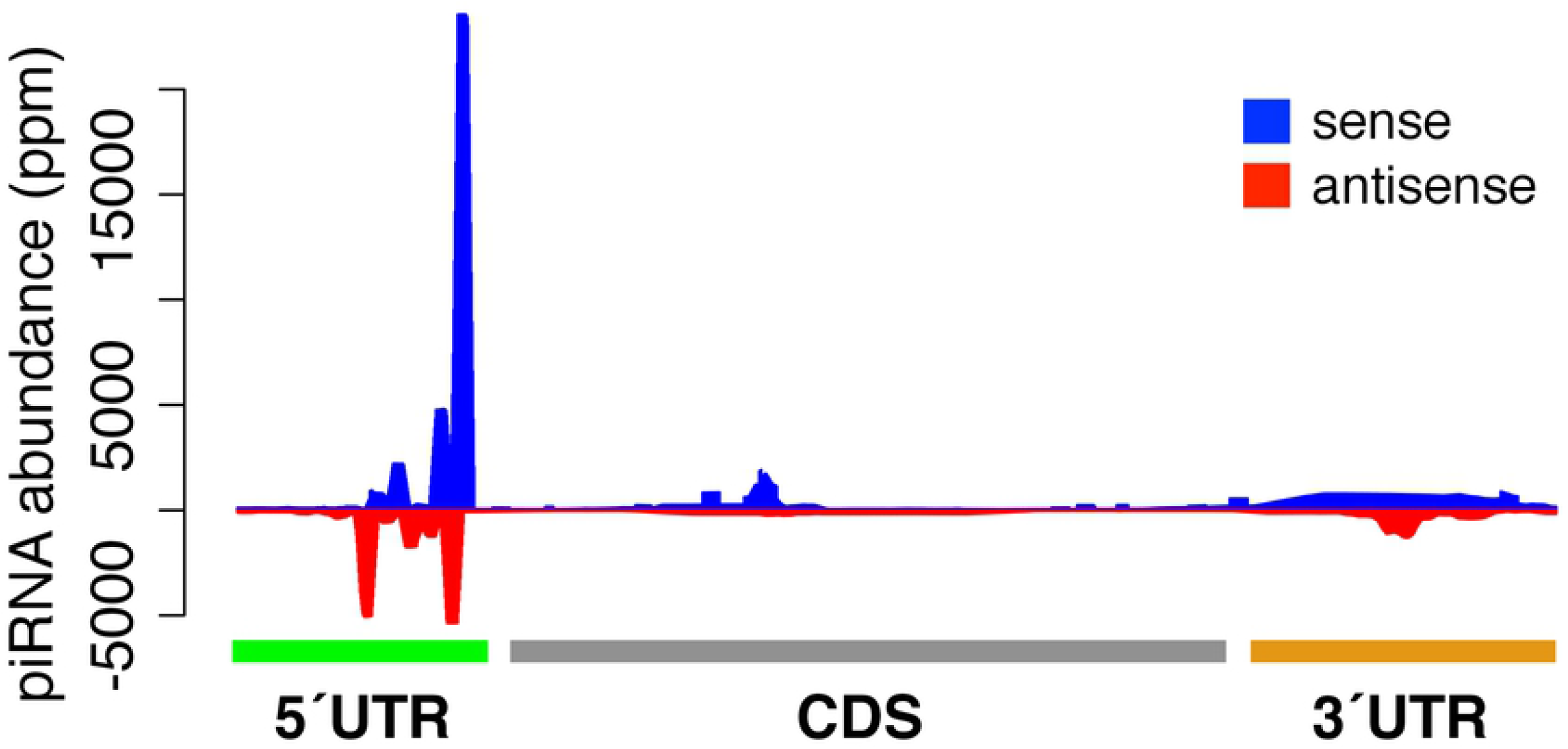
Relative mapping positions of genic piRNAs. The relative mapping position of piRNAs on highly expressed genes. piRNAs mapping to the sense and antisense strands of genes are highlighted in blue and red respectively.

### piRNA biogenesis pathway factors are differentially expressed during ISM atrophy and the commitment to die

We next sought to determine which piRNA pathway protein components are expressed in the ISMs. Based on piRNA pathway genes characterized in *Drosophila,* we identified all 20 of the genes we sought in the ISMs *(ago3, armi, aub, piwi, BoYo, egg, krimp, mael, papi, qin, rhi, shu, spn-E, tej, tud, vas, vret, Tejas, Hen-1,* BoYB, and *zuc),* plus two small RNA biogenesis pathway factors (*ago1* and *ago2*) In agreement with published reports, we only identified a single Aub/Piwi protein sequence, which is also the case for other Lepidopterans like *Tricoplisia ni* and *Bombyx mori* (55). Consequently, we refer to this protein as aub/piwi.

Next, we examined the mRNA-seq reads from six developmental stages (days 13, 14, 15, 16, 17, 18) plus 20E-treated to determine if these factors are differentially expressed. While the expression levels of *ago3* and *aub* appear to be low, their expression was nevertheless ranked within the top 64.4% and 31.1% respectively of all expressed genes (Figure 4A). The expression levels of other genes in this pathway were also within the top 18%-65% of all expressed genes (Figure S4).

**Figure 4:**
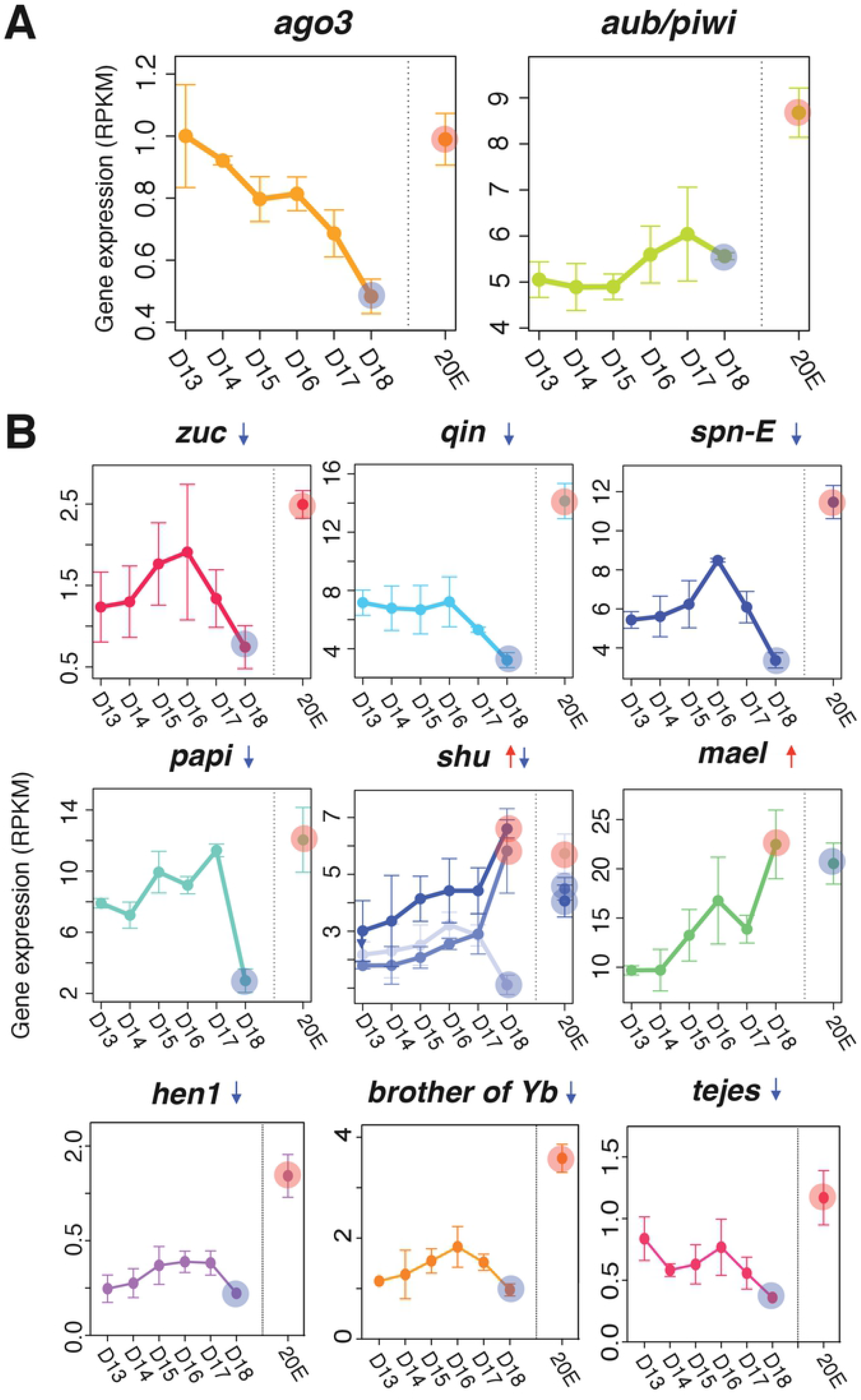
Expression of piRNA biogenesis pathway factors. **A**: Gene expression of *ago3* and *aub/piwi* based on RNA-seq analyses (top) **B**: Downregulated (*zuc, qin, spn-E, hen1, brother of Yb, tejes* and *papi*), mixed (*shu*), and upregulated (*mael*) piRNA pathway factors in the ISMs prior to the initiation of cell death. Red and blue circles on day 18 indicate up- and down-regulation of the factors compared to expression on day 13. Those circles in the 20E lane indicate statistically significant changes in ISM gene expression animals injected on day 17 with 20E to delay cell death on day 18.

Computational analysis suggests that the expression of many of the small RNA biogenesis pathway factors were developmentally regulated, with a general trend for stable or increasing expression prior to the initiation of atrophy (days 13-16), a sharp decline on day 17, and a near loss of expression when the ISMs became commitment to die on day 18. Four genes were significantly repressed on day 18:*papi (q* = 5.8 × 10^−9^, fc = −2.6), *qin* (q = 2.9 × 10^−7^, fc = −2.1), *zuc* (q = 3.9 × 10^−3^, fc = −2.1), and *spn-E* (q = 1.7 × 10^−9^, fc = −2.0) (Figure 4B) and five other demonstrated this general trend (*ago3*, *armi*, *egg*, *krimp*, and *tud*), although their changes did not reach statistical significance (Figure S4). In contrast, *mael* mRNA levels were increased with the commitment to die (q = 6.4 × 10^−19^, fc = 2.5; Figure 4B) as were some members of the *shu* family (Figure 4B). In all cases, expression of piRNA pathway components were regulated by 20E and displayed their highest levels of expression when exposed to the exogenous steroid, suggesting that these genes are regulated by the same developmental signals that control ISM atrophy and death.

### Transposon expression becomes deregulated when the ISM become committed to die

We observed that the abundance of both piRNAs, and the majority of the factors that mediate their biogenesis, declined precipitously when the ISMs became committed to die on day 18. Consequently, we sought to test the hypothesis that this loss might lead to changes in transposon expression. Using the RNA-seq reads mapped to transposon loci as the input, we computed the expression of each transposable element and the fold change between pairs of stages during: homeostasis (day 13 vs day 15); atrophy (day 14 vs day 17) and commitment to die (day 17 and day 18); as well as the response to hormone treatment (day 18 vs 20E) (Figure 5). There were no significant differences in transposon expression between the pairs of homeostatic or atrophic muscles. Only the Tourist DNA mobile element was up-regulated on day 17 (q = 7.0 × 10^−7^, fc = 2.1) compared to day 13. Once the muscle became committed to die (but prior to the initiation of death later in the day), the patterns of the transposon expression changed dramatically. Four DNA transposable elements were up-regulated compared to those on day 13: Tourist (*q* = 8.0 × 10^−13^, fc = 4.0), hAT-Pegasus (*q* = 4.1 × 10^−5^, fc = 2.6), TcMar-Tc1 (*q* = 6.4 × 10^−12^, fc = 2.2), and general DNA mobile elements (*q* = 4.6 × 10^−11^, fc = 2.4). Concurrently, the expression levels of four transposons were significantly down-regulated on day 18 as compared to day 13: 5S-Deu (SINE; *q* = 4.7 × 10^−4^, fc = −2.0), LOA (LINE; *q* = 1.1 × 10^−8^, fc = −2.0), Penelope (LINE; *q* = 3.1 × 10^−6^, fc = −2.6), and CMC-Transib (DNA transposon; *q* = 4.7 × 10^−4^, fc = −2.3). Interestingly, when ISM death was delayed with steroid injection (20E), the differential expression for most of the developmentally-regulated transposons was muted.

**Figure 5:**
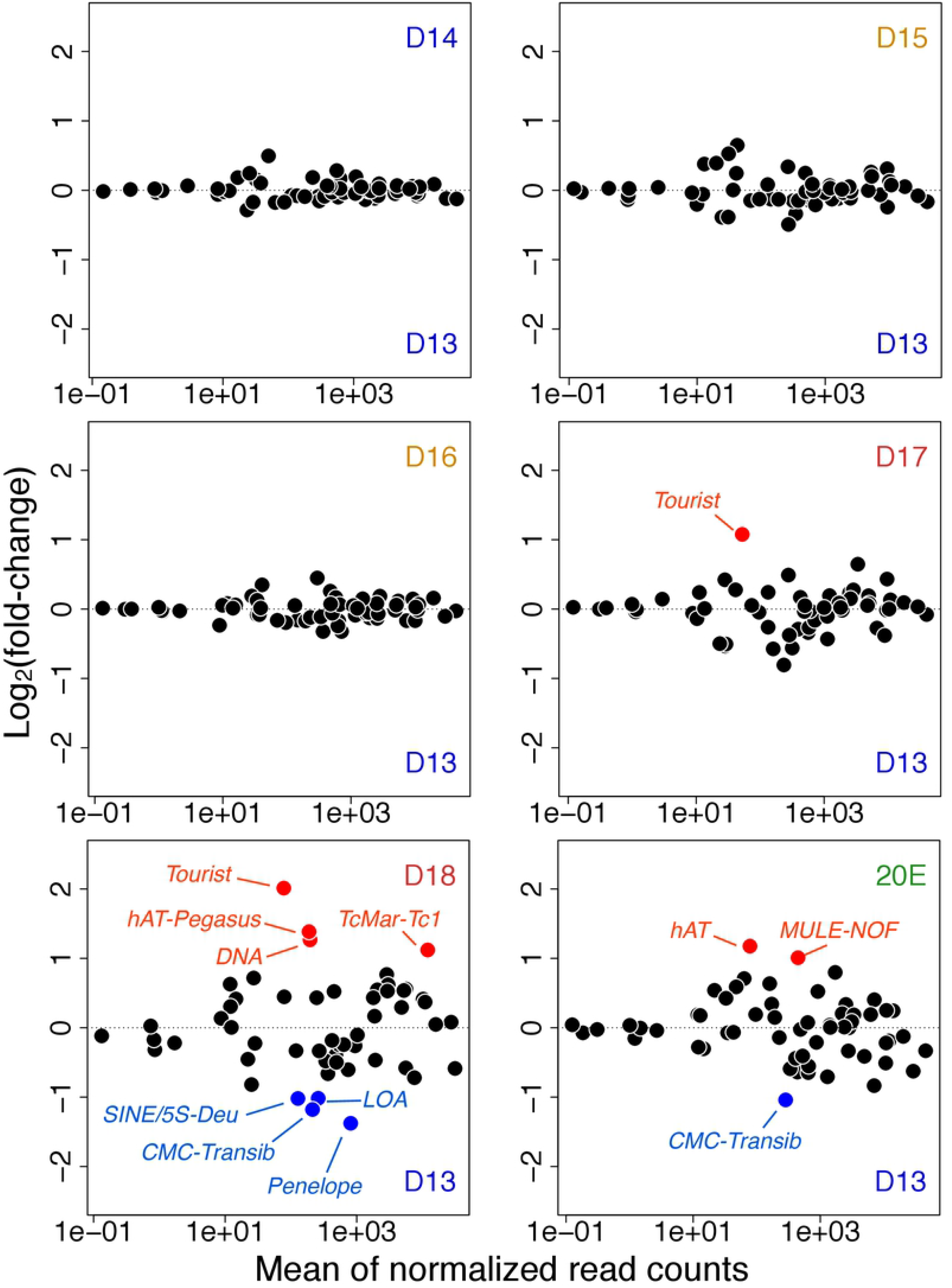
Transposon expression in atrophy and ping-pong piRNAs. Changes in transposon expression during ISM development relative to day 13 (from day 14 to day 18, and 20E). Up-regulated and down-regulated transposons are highlighted in red and blue respectively.

To help understand the molecular mechanisms that could facilitate differential transposon expression, we computed the correlation between transposon expression and ping-pong Z-score for the associated piRNAs (Figure S5). There was a general inverse relationship between the induced transposons and their corresponding piRNAs for the Tourist, hAT-Pegasus, and DNA element families (except for TcMar-Tc1). There were three families where there was no clear relationship between piRNA and transposon abundance. In the case of the down-regulated transposon families, only SINE/5S-Deu retrotransposons displayed the loss of associated piRNAs. As a general observation, transposon expression decreased when piRNAs were ping-pong amplified on the same or previous day (e.g. those of DNA/Kolobok-Hydra, DNA/P, DNA/hAT-Ac, and LINE/Jockey in Figure S5). Taken together, these data suggest that there are significant changes in transposon expression when the ISMs become committed to die that may be mediated by corresponding piRNA abundance.

## Discussion

In this report we present substantial evidence supporting the presence of developmentally-regulated piRNAs in striated skeletal muscle, a highly differentiated somatic cell. Like piRNAs characterized from ovary and testis, *Manduca* ISM piRNAs: are ~27 nucleotides long, have a strong 5’ uridine bias, are oxidation resistant, amplify via a ping-pong mechanism, map to transposable elements, and are predominantly antisense. (Efforts to demonstrate that these piRNAs were physically bound to Aub were unsuccessful as none of the anti-*Drosophila* Aub antibodies that we tested recognized or precipitated *Manduca* Aub/Piwi (data not shown)). These piRNAs represent a substantial percentage of the small RNA pool within the tissue and are neither low abundance sequences nor potential contaminants from mRNA degradation. The ISMs are a very large and discrete tissue that can be isolated cleanly from the animal without contamination from other piRNA-rich tissues such as gonads (39). As well, unlike mammalian muscle, the ISMs are composed of only a single fiber type and do not contain regenerative stem cells like satellite cells or pericytes that might complicate the analysis (56).

The expression of piRNAs and the machinery required for their synthesis and action in somatic cells is controversial and the subject of debate (24, 27, 29). piRNAs have been also reported in a small number of somatic cells such as cancer cells (Mei et al., 2013) and regenerative stem-like cells in invertebrates (57). For example, PIWI-like proteins and piRNAs have been identified in somatic cells of the Cnidarian *Hydra* and the flatworm *Macrostomum,* although it is thought that their expression is restricted to stem-like progenitor cells that have the capacity to regenerate all cell types of the animal including gonad (58, 59). Loss of somatic piRNA pathway in *Drosophila* fat body disrupts metabolic homeostasis by depleting lipids and stored metabolites leading shortened lifespan (54). In the silkmoth *Bombyx mori*, the primary sex-determining factor is regulated by a single piRNA originated from the sex-determining genomic locus of W chromosome (60). In the nematode *C. elegans,* the piRNA pathway components, including PRDE-1 and PRG-1, are expressed in neurons and their inhibition facilitates sensory neuron regeneration following injury in a cell autonomous manner (61).

The best characterized system for the analysis of somatic piRNAs is follicle cells in the *Drosophila* ovary (27, 62, 63). Follicle cells are epithelial cells that are needed for the survival and maturation of the underlying germline cells, and disruption of the piRNA pathway in these cells leads to enhanced transposon activity and sterility (62). While follicle cells are derived somatically, they are physically connected to the germline via large ring canals, so it is not clear how much interplay may exist between them. Nevertheless, piRNA production can take place in the follicle-derived OSS (ovary somatic sheet) cell line, supporting the hypothesis that these specialized somatic cells can produce piRNAs (22, 51, 64, 65). However, piRNA production in these cells differs from that seen in ovarian cells in several key ways (14, 51). First, follicle cells lack both Aub and Ago3, and consequently do not display ping-pong amplification (14). Secondly, they preferentially express piRNAs from uni-strand genomic piRNA clusters, while ovarian cells produce piRNAs from both single and double-stranded transposable element clusters (14, 66). In contrast to follicle cells, the ISMs express both *Aub* and *Ago3* and display efficient ping-pong amplification with high Z-scores. These data suggest that while the ISMs are clearly somatic cells, their piRNA pathway more closely represents the hallmarks of germline piRNA production.

Sequencing and mapping analyses have demonstrated that *Manduca* piRNAs are generated from both transposable elements and protein coding genes. In the few instances where genic piRNAs have been analyzed, they have been shown to be derived predominantly from the 3’ UTR sequences within mRNAs (52–54, 64). In contrast, genic piRNAs analyzed from *Manduca* appear to be derived primarily from the start of translation in the 5’UTR. The possible regulatory role that these sequences might serve is unclear. There is the intriguing observation that mRNAs become more labile in day 17 ISMs, which may facilitate the rapid upregulation of death-associated transcripts (47). It is not known if these genic piRNAs might participate in this process via some uncharacterized mechanism. However, considering the instability of ISM transcript mediated by 3’ UTRs, we speculated that miRNAs could play a role in facilitating the rapid transition from the atrophy program to the one that mediates death. Indeed, we have demonstrated that miRNAs targeting the 3’UTRs of death-associated transcripts can repress translation (42).

The ISMs are a classic model system for the study of skeletal muscle atrophy and death (67–69). Data presented here demonstrate that the expression of ISM piRNAs is developmentally-regulated, with low levels during atrophy and peak expression on day 17 in advance of the commitment of the muscles to die late on day 18. Consequently, we examined the expression of transposable elements since they are the primary target of piRNAs. Transposon expression appears to be tightly controlled in the ISMs since there was almost no variation their abundance until day 17, at which point they became deregulated. Within a matter of hours, the expression of several DNA transposons increased significantly, while some elements, most notably retrotransposons, were concurrently repressed. The ability of the ISMs to initiate PCD occurs when the circulating levels of the steroid hormone 20E decline below a specific threshold on day 17 (44). Hormone replacement with exogenous 20E on day 17 not only delays ISM death, it also prevents the developmental changes in piRNA and transposon expression that accompany PCD. This data supports the hypothesis that this pathway is also under hormonal control. It should be noted that the changes in transposon expression occur well in advance of the initiation of cell death and therefore do not appear to be a secondary consequence of cellular suicide. In the germline, transposon expression is repressed in order to protect the genome from insertional mutagenesis and subsequent catastrophe. However, this same process may confer an advantage for the organism by ensuring that cell death is indeed an irreversible process. For example, during apoptosis, genome destruction is insured by the activation of endogenous nucleases that cleave chromosomal DNA into nucleosome sized fragments (70). However, the ISMs die by an autophagic process that does not include DNA fragmentation (56, 67, 71). Perhaps transposon expression and reintegration serve a similar role for cells undergoing non-apoptotic forms of cell death where it insures that condemned cells are truly “dead” by fragmenting the genome and depleting the cell of beta-nicotinamide adenine dinucleotide and ATP (72). Some support for this hypothesis comes from the observation that transposon expression is elevated in certain neurodegenerative disorders (73–75), a phenomenon that has been called a “transposon storm” (76, 77). To date, none of these studies have examined the possible expression of piRNAs in the diseased tissues. We attempted to directly test the hypothesis that reintegration of transposons in *Manduca* would result in double stranded breaks in genomic DNA by sequencing genomic libraries generated from ISMs taken before and after adult eclosion. Unfortunately, laboratory reared *Manduca* are not isogenic, and the individual-to-individual variability in transposon copy number precluded a direct test of the hypothesis (data not shown). As well, we tried to use antibodies against phosphorylated histone H2Av since it is a marker of double stranded DNA breaks (78), but the antibodies directed against *Drosophila* phosphorylated H2Av did not cross react with the *Manduca* protein, and a Lepidopteran-specific antibody has not been identified (data not shown).

Taken together, we have presented compelling evidence that piRNAs are expressed in highly differentiated somatic cells. Their expression is developmentally regulated by the steroid hormone 20E. piRNA expression is repressed when the ISMs become committed to die, which is correlated with the deregulation of transposon expression. The expression and possible re-integration of transposons may help insure that the muscles, which do not die by apoptosis, nevertheless experience genome degradation. The work both verifies the expression of somatic piRNAs and may provide new insights into degenerative processes during aging and pathogenesis.

## Materials and Methods

### Animals

The tobacco hawkmoth *Manduca sexta* was reared and staged as described previously (44). The lateral intersegmental muscles (ISMs) were dissected free from adjacent tissue under ice-cold saline, flash frozen on dry ice and stored in liquid nitrogen until used for RNA isolation.

Some animals were injected on day 17 of pupal-adult development with 25ug of 20-hydroxyecdysone (20E) (Sigma) in 10% isopropanol to delay ISM death (79) and then the ISMs removed prior to the normal time of eclosion on day 18.

### RNA Isolation, Library Construction and Sequencing

The ISMs of 3-4 animals per developmental time point (eight time points in total: days 13, 14, 15, 16, 17, 18, and 1-hour post-eclosion (PE); plus 20-hydroxyecdysone injection on day 17: 20E) were homogenized and total RNA was isolated using a mirVana RNA Isolation kit (Life Technologies).

For small RNA-seq library construction, 50 ug of total RNA was fractionated by 15% urea polyacrylamide gel electrophoresis and the 18-30 nt fraction extracted for library construction. 3’ and 5’ adaptors were ligated to the small RNA and the cDNA reverse transcribed and PCR amplified. The libraries were purified by polyacrylamide gel electrophoresis, and subjected to 50 nt single-end sequencing on an Illumina HiSeq™ 2000 (San Diego, CA) by Beijing Genomics Institute (Hong Kong).

For RNA-seq, directional and random primed cDNA libraries were constructed with poly(A)+ RNA, analyzed with a Bioanalyzer (Agilent Technologies; Santa Clara, CA) and 50 nt single-end sequencing was performed as above. For each of the eight time points, we prepared three biological replicates of RNA-seq libraries.

All the sequencing libraries are accessible from Gene Expression Omnibus (GEO) (accession number GSE80830)

### Oxidized small RNA-seq library

Unlike miRNAs, piRNAs are 2’-O-methylated at their 3’ termini, which renders them resistant to oxidation (48). Therefore, we oxidized small RNAs as outlined in (16), then cloned the resulting piRNAs as above. We sequenced the oxidized small RNA library at the Deep Sequencing Core Facility at the UMass Medical School.

### Genomic sequence and annotation data

We downloaded the genomic assembly of *Manduca sexta* (Msex1.0) and the transcript and the protein sequences (revised-OGS-June2012) from the *Manduca* Base (http://agripestbase.org/manduca/) (42). The genomic assembly contains 20,868 scaffolds, with a median length of 994 bp. We annotated protein-coding genes, transposable elements, low complexity regions, miRNAs, and other non-coding RNAs such as rRNA, tRNA, snoRNA, snRNA etc. The detailed annotation protocol and statistics are described in Supplementary materials (below). All the sequencing libraries are accessible from GSE80830 in Gene Expression Omnibus.

### Sequence extraction and annotation of piRNAs

After computationally removing the adaptor sequences, we mapped the extracted sequences to the reference *Manduca* genome using the Bowtie algorithm (80). We only retained reads that matched the genome perfectly for downstream analysis. To identify potential piRNAs, we selected sequences that were longer than 22 nt and not annotated as miRNAs or other non-coding RNAs (see Supplementary Materials for annotation of miRNAs and other non-coding RNAs). The reads of piRNAs that mapped to multiple loci in the genome were apportioned over these loci, and piRNA reads were normalized by the total number of genome mapping reads excluded other non-coding RNAs (rRNA, tRNA etc.). piRNA abundance is quantified in parts per million (ppm).

### piRNA ping-pong signature

To determine if *Manduca* piRNAs are amplified via the ping-pong cycle, we computed the Z-score as described in (51). Briefly, we identified the piRNA pairs that mapped to overlapping genomic positions but on opposite genomic strands. We counted the numbers of such pairs with 5’-5’ overlapping distances from 1 to 20 nts, and calculated Z-score for the 10-nt overlap (the expected overlapping distance due to ping-pong) using the counts of 1-9 nt and 11-19 nt overlaps as the background.

### Relative mapping position of piRNAs on genes

In order to characterize the relative positions in mRNAs that *Manduca* piRNAs map to, we scaled the 5’ UTRs, coding regions, and 3’ UTRs of mRNAs to 350, 1000, and 800 nts respectively, which we calculated are the median lengths of these regions in annotated *M. sexta* genes. Using scaled non-overlapping windows which are equivalent to each of the 2,150 nts of the scaled genes, we counted piRNA abundance in RPKM.

### Gene expression and differentially expressed genes

We mapped reads in each RNA-seq library to the reference *Manduca* genome using the TopHat2 algorithm (81) allowing 2 mismatches (“-v 2”). To detect reads mapping to transposons, we allowed reads to map to multiple locations of the genome with the “-g” option in TopHat2. Since the most abundant transposon in *Manduca* is SINE, with 44,487 copies when all subfamilies are combined (Table S2), we ran TopHat2 with “-g 45,000”. Reads were apportioned by the number of times they mapped to the genome.

After mapping, we counted the number of RNA-seq reads for each gene and transposon, expressed in the unit of RPKM (Reads Per Kilobase of transcript per Million mapped reads). To identify differentially expressed genes and transposons between a pair of time points during ISM development, we ran the DESeq2 algorithm (version 1.5.5) implemented in R (Anders and Huber, 2010) using mapped read counts as the input, taking advantage of the three biological replicates per stage. Genes with q-value < 0.01 and absolute fold-change (fc) > 2 were considered to be differentially expressed between the two time points.

## Acknowledgements

We thank Dr. Wendy Smith for the provision of animals and Dr. Leonid Pobezinsky for helpful discussions. This work was supported by a Life Sciences Moment Fund grant from the University of Massachusetts Center for Clinical and Translational Science to LMS, ZW and WT, the Eugene M. and Ronnie Isenberg Professorship Endowment (LMS), NIH grant P01HD078253 to WT and ZW, and National Institute of Child Health and Human Development Grant P01HD078253 to W. E. Theurkauf and Z. Weng.

## Supporting information

**S1 Methods:** Methods for genomic annotation

**S1 Table:** Small RNA-seq reads statistics.

**S2 Table:** Transposable elements in *M. sexta.*

**S1 Figure:** Length distribution, sequence logos, and genomic annotations of small RNAs in the ISM development.

**S2 Figure:** Expression and change of piRNAs mapped to transposon families in the ISM development.

**S3 Figure:** piRNA relative mapping positions on genes.

**S4 Figure:** Gene expression of small RNA biogenesis pathway factors.

**S5 Figure:** Transposon expression and ping-pong piRNA Z-scores.

## Data reporting

All the sequencing libraries are accessible from GSE80830 in Gene Expression Omnibus.

## Supplemental Figures

**Figure S1:** Length distribution, sequence logos, and genomic annotations of small RNAs in the ISM development. Bar plots in the leftmost column display the length distribution of small RNAs in the ISM developmental stages. Pie charts in the second left column show genomic annotations of small RNAs in the *M. sexta* genome. Two sequence logos for miRNAs and reads longer than 23nt are shown for each developmental stage. Pie charts in the rightmost column show genomic annotations of reads longer than 23nt. Percentages without parentheses represent relative abundance in the reads longer than 23nt, and percentages within parentheses represent the abundance in total reads.

**Figure S2:** Expression and change of piRNAs mapped to transposon families in the ISM development. The expression of piRNAs mapped to each transposon family is shown. Shaded bars indicate the fractions of piRNAs amplified via the ping-pong amplification. The numbers in the parentheses are copy numbers of transposon families in the *M. sexta* genome.

**Figure S3:** piRNA relative mapping positions on genes. The distribution of piRNA abundance on highly and low expressed genes in each time point is shown. piRNAs mapped to sense and antisense are highlighted in blue and red respectively.

**Figure S4**: Gene expression of small RNA biogenesis pathway factors. Red and blue circles on day 18 indicate up- and down-regulation of the factors compared to those on day 13. Similarly, those circles on 20E indicate up- and down-regulation of the factors compared to those on day 18. Only a single *aub/piwi* gene (Msex009073) is detected in the genome.

**Figure S5:** Transposon expression and ping-pong piRNA Z-scores. Left Y-axis indicates transposon expression in RPKM, and right Y-axis indicates ping-pong Z-scores of the piRNAs mapped to the transposon. Red and blue diamonds show up- and down-regulation of transposon expression. Statistically significant Z-scores are colored in light blue, while non-significant ones are grey. The numbers in the parentheses are copy numbers of transposon families in the *M. sexta* genome.

## Supplemental Tables

**Table S1: Small RNA-seq reads statistics.** The sample on day 13 has both unoxidized and oxidized small RNA-seq libraries. “ncRNAs” refer to non-coding RNAs such as tRNA, rRNA, and snoRNA. “Reads excluding ncRNAs and miRNAs” correspond to the piRNAs analyzed in this study. “Transposon matching reads” and “Gene matching reads” indicate piRNAs mapped to transposons and genes respectively. Due to the fact that a piRNA read can map to both the sense and the antisense orientations of a transposon, the sum of these transposon matching reads is greater than the total number of transposon matching reads. The numbers in parentheses avoid this discrepancy by apportioning a value of 0.5 to sense and antisense for each read that maps to both orientations.

**Table S2: Transposable elements in *M. sexta.*** Copy numbers, total bases, and fractions within the *M. sexta* genome are shown for the 64 transposon families detected with RepeatMasker.

## References

1. Castel SE, Martienssen RA. RNA interference in the nucleus: roles for small RNAs in transcription, epigenetics and beyond. Nat Rev Genet. 2013;14(2):100–12.

2. Ghildiyal M, Zamore PD. Small silencing RNAs: an expanding universe. Nat Rev Genet. 2009;10(2):94–108.

3. Grewal SI, Elgin SC. Transcription and RNA interference in the formation of heterochromatin. Nature. 2007;447(7143):399–406.

4. Bartel DP. Metazoan MicroRNAs. Cell. 2018;173(1):20–51.

5. Guo H, Ingolia NT, Weissman JS, Bartel DP. Mammalian microRNAs predominantly act to decrease target mRNA levels. Nature. 2010;466(7308):835–40.

6. Ha M, Kim VN. Regulation of microRNA biogenesis. Nat Rev Mol Cell Biol. 2014;15(8):509–24.

7. Czech B, Hannon GJ. One Loop to Rule Them All: The Ping-Pong Cycle and piRNA-Guided Silencing. Trends Biochem Sci. 2016;41(4):324–37.

8. Czech B, Munafo M, Ciabrelli F, Eastwood EL, Fabry MH, Kneuss E, et al. piRNA-Guided Genome Defense: From Biogenesis to Silencing. Annu Rev Genet. 2018;52:131–57.

9. Iwasaki YW, Siomi MC, Siomi H. PIWI-Interacting RNA: Its Biogenesis and Functions. Annu Rev Biochem. 2015;84:405–33.

10. Williams Z, Morozov P, Mihailovic A, Lin C, Puvvula PK, Juranek S, et al. Discovery and Characterization of piRNAs in the Human Fetal Ovary. Cell Rep. 2015;13(4):854–63.

11. Ozata DM, Gainetdinov I, Zoch A, O’Carroll D, Zamore PD. PIWI-interacting RNAs: small RNAs with big functions. Nat Rev Genet. 2019;20(2):89–108.

12. Huang X, Fejes Toth K, Aravin AA. piRNA Biogenesis in Drosophila melanogaster. Trends Genet. 2017;33(11):882–94.

13. Brennecke J, Aravin AA, Stark A, Dus M, Kellis M, Sachidanandam R, et al. Discrete small RNA-generating loci as master regulators of transposon activity in Drosophila. Cell. 2007;128(6):1089–103.

14. Malone CD, Brennecke J, Dus M, Stark A, McCombie WR, Sachidanandam R, et al. Specialized piRNA pathways act in germline and somatic tissues of the Drosophila ovary. Cell. 2009;137(3):522–35.

15. Thomson T, Lin H. The biogenesis and function of PIWI proteins and piRNAs: progress and prospect. Annu Rev Cell Dev Biol. 2009;25:355–76.

16. Gunawardane LS, Saito K, Nishida KM, Miyoshi K, Kawamura Y, Nagami T, et al. A slicer-mediated mechanism for repeat-associated siRNA 5’ end formation in Drosophila. Science. 2007;315(5818):1587–90.

17. Cora E, Pandey RR, Xiol J, Taylor J, Sachidanandam R, McCarthy AA, et al. The MID-PIWI module of Piwi proteins specifies nucleotide-and strand-biases of piRNAs. RNA. 2014;20(6):773–81.

18. Stein CB, Genzor P, Mitra S, Elchert AR, Ipsaro JJ, Benner L, et al. Decoding the 5’ nucleotide bias of PIWI-interacting RNAs. Nat Commun. 2019;10(1):828.

19. Wang W, Yoshikawa M, Han BW, Izumi N, Tomari Y, Weng Z, et al. The initial uridine of primary piRNAs does not create the tenth adenine that Is the hallmark of secondary piRNAs. Mol Cell. 2014;56(5):708–16.

20. Klattenhoff C, Theurkauf W. Biogenesis and germline functions of piRNAs. Development. 2008;135(1):3–9.

21. Weick EM, Miska EA. piRNAs: from biogenesis to function. Development. 2014;141(18):3458–71.

22. Sato K, Siomi MC. The piRNA pathway in Drosophila ovarian germ and somatic cells. Proc Jpn Acad Ser B Phys Biol Sci. 2020;96(1):32–42.

23. Rojas-Rios P, Simonelig M. piRNAs and PIWI proteins: regulators of gene expression in development and stem cells. Development. 2018;145(17).

24. Pandey RR, Homolka D, Chen KM, Sachidanandam R, Fauvarque MO, Pillai RS. Recruitment of Armitage and Yb to a transcript triggers its phased processing into primary piRNAs in Drosophila ovaries. PLoS Genet. 2017;13(8):e1006956.

25. Perera BPU, Tsai ZT, Colwell ML, Jones TR, Goodrich JM, Wang K, et al. Somatic expression of piRNA and associated machinery in the mouse identifies short, tissue-specific piRNA. Epigenetics. 2019;14(5):504–21.

26. Teefy BB, Siebert S, Cazet JF, Lin H, Juliano CE. PIWI-piRNA pathway-mediated transposable element repression in Hydra somatic stem cells. RNA. 2020;26(5):550–63.

27. Onishi R, Sato K, Murano K, Negishi L, Siomi H, Siomi MC. Piwi suppresses transcription of Brahma-dependent transposons via Maelstrom in ovarian somatic cells. Sci Adv. 2020;6(50).

28. Huang S, Ichikawa Y, Igarashi Y, Yoshitake K, Kinoshita S, Omori F, et al. Piwi-interacting RNA (piRNA) expression patterns in pearl oyster (Pinctada fucata) somatic tissues. Sci Rep. 2019;9(1):247.

29. Lewis SH, Quarles KA, Yang Y, Tanguy M, Frezal L, Smith SA, et al. Pan-arthropod analysis reveals somatic piRNAs as an ancestral defence against transposable elements. Nat Ecol Evol. 2018;2(1):174–81.

30. Feng M, Kolliopoulou A, Zhou YH, Fei SG, Xia JM, Swevers L, et al. The piRNA response to BmNPV infection in the silkworm fat body and midgut. Insect Sci. 2020.

31. Whitfield ZJ, Dolan PT, Kunitomi M, Tassetto M, Seetin MG, Oh S, et al. The Diversity, Structure, and Function of Heritable Adaptive Immunity Sequences in the Aedes aegypti Genome. Curr Biol. 2017;27(22):3511–9 e7.

32. Tassetto M, Kunitomi M, Whitfield ZJ, Dolan PT, Sanchez-Vargas I, Garcia-Knight M, et al. Control of RNA viruses in mosquito cells through the acquisition of vDNA and endogenous viral elements. Elife. 2019;8.

33. Perrat PN, DasGupta S, Wang J, Theurkauf W, Weng Z, Rosbash M, et al. Transposition-driven genomic heterogeneity in the Drosophila brain. Science. 2013;340(6128):91–5.

34. Tindell SJ, Rouchka EC, Arkov AL. Glial granules contain germline proteins in the Drosophila brain, which regulate brain transcriptome. Commun Biol. 2020;3(1):699.

35. Lee EJ, Banerjee S, Zhou H, Jammalamadaka A, Arcila M, Manjunath BS, et al. Identification of piRNAs in the central nervous system. RNA. 2011;17(6):1090–9.

36. Rajasethupathy P, Antonov I, Sheridan R, Frey S, Sander C, Tuschl T, et al. A role for neuronal piRNAs in the epigenetic control of memory-related synaptic plasticity. Cell. 2012;149(3):693–707.

37. Sheel A, Shao R, Brown C, Johnson J, Hamilton A, Sun D, et al. Acheron/Larp6 Is a Survival Protein That Protects Skeletal Muscle From Programmed Cell Death During Development. Front Cell Dev Biol. 2020;8:622.

38. Schwartz LM, Truman JW. Peptide and steroid regulation of muscle degeneration in an insect. Science. 1982;215(4538):1420–1.

39. Schwartz LM. Skeletal Muscles Do Not Undergo Apoptosis During Either Atrophy or Programmed Cell Death-Revisiting the Myonuclear Domain Hypothesis. Front Physiol. 2018;9:1887.

40. Schwartz LM, Ruff RL. Changes in contractile properties of skeletal muscle during developmentally programmed atrophy and death. Am J Physiol Cell Physiol. 2002;282(6):C1270–7.

41. Lockshin RA, Williams CM. Programmed Cell Death--I. Cytology of Degeneration in the Intersegmental Muscles of the Pernyi Silkmoth. J Insect Physiol. 1965;11:123–33.

42. Tsuji J, Thomson T, Chan E, Brown CK, Oppenheimer J, Bigelow C, et al. High-resolution analysis of differential gene expression during skeletal muscle atrophy and programmed cell death. Physiol Genomics. 2020;52(10):492–511.

43. Jones ME, Schwartz LM. Not all muscles meet the same fate when they die. Cell Biol Int. 2001;25(6):539–45.

44. Schwartz LM, Truman JW. Hormonal control of rates of metamorphic development in the tobacco hornworm Manduca sexta. Dev Biol. 1983;99(1):103–14.

45. Schwartz LM, Kosz L, Kay BK. Gene activation is required for developmentally programmed cell death. Proc Natl Acad Sci U S A. 1990;87(17):6594–8.

46. Schwartz LM, Myer A, Kosz L, Engelstein M, Maier C. Activation of polyubiquitin gene expression during developmentally programmed cell death. Neuron. 1990;5(4):411–9.

47. Cascone PJ, Schwartz LM. Post-transcriptional regulation of gene expression during the programmed death of insect skeletal muscle. Dev Genes Evol. 2001;211(8-9):397–405.

48. Vagin VV, Sigova A, Li C, Seitz H, Gvozdev V, Zamore PD. A distinct small RNA pathway silences selfish genetic elements in the germline. Science. 2006;313(5785):320–4.

49. Zhang Z, Xu J, Koppetsch BS, Wang J, Tipping C, Ma S, et al. Heterotypic piRNA Ping-Pong requires qin, a protein with both E3 ligase and Tudor domains. Mol Cell. 2011;44(4):572–84.

50. Meister G. Argonaute proteins: functional insights and emerging roles. Nat Rev Genet. 2013;14(7):447–59.

51. Li C, Vagin VV, Lee S, Xu J, Ma S, Xi H, et al. Collapse of germline piRNAs in the absence of Argonaute3 reveals somatic piRNAs in flies. Cell. 2009;137(3):509–21.

52. Robine N, Lau NC, Balla S, Jin Z, Okamura K, Kuramochi-Miyagawa S, et al. A broadly conserved pathway generates 3’UTR-directed primary piRNAs. Curr Biol. 2009;19(24):2066–76.

53. Sokolova OA, Ilyin AA, Poltavets AS, Nenasheva VV, Mikhaleva EA, Shevelyov YY, et al. Yb body assembly on the flamenco piRNA precursor transcripts reduces genic piRNA production. Mol Biol Cell. 2019;30(12):1544–54.

54. Jones BC, Wood JG, Chang C, Tam AD, Franklin MJ, Siegel ER, et al. A somatic piRNA pathway in the Drosophila fat body ensures metabolic homeostasis and normal lifespan. Nat Commun. 2016;7:13856.

55. Cao X, He Y, Hu Y, Wang Y, Chen YR, Bryant B, et al. The immune signaling pathways of Manduca sexta. Insect Biochem Mol Biol. 2015;62:64–74.

56. Beaulaton J, Lockshin RA. Ultrastructural study of the normal degeneration of the intersegmental muscles of Anthereae polyphemus and Manduca sexta (Insecta, Lepidoptera) with particular reference of cellular autophagy. J Morphol. 1977;154(1):39–57.

57. Kim IV, Riedelbauch S, Kuhn CD. The piRNA pathway in planarian flatworms: new model, new insights. Biol Chem. 2020;401(10):1123–41.

58. Juliano CE, Reich A, Liu N, Gotzfried J, Zhong M, Uman S, et al. PIWI proteins and PIWI-interacting RNAs function in Hydra somatic stem cells. Proc Natl Acad Sci U S A. 2014;111(1):337–42.

59. Zhou X, Battistoni G, El Demerdash O, Gurtowski J, Wunderer J, Falciatori I, et al. Dual functions of Macpiwi1 in transposon silencing and stem cell maintenance in the flatworm Macrostomum lignano. RNA. 2015;21(11):1885–97.

60. Kiuchi T, Koga H, Kawamoto M, Shoji K, Sakai H, Arai Y, et al. A single female-specific piRNA is the primary determiner of sex in the silkworm. Nature. 2014;509(7502):633–6.

61. Kim KW, Tang NH, Andrusiak MG, Wu Z, Chisholm AD, Jin Y. A Neuronal piRNA Pathway Inhibits Axon Regeneration in C. elegans. Neuron. 2018;97(3):511–9 e6.

62. Olivieri D, Sykora MM, Sachidanandam R, Mechtler K, Brennecke J. An in vivo RNAi assay identifies major genetic and cellular requirements for primary piRNA biogenesis in Drosophila. EMBO J. 2010;29(19):3301–17.

63. Ishizu H, Iwasaki YW, Hirakata S, Ozaki H, Iwasaki W, Siomi H, et al. Somatic Primary piRNA Biogenesis Driven by cis-Acting RNA Elements and trans-Acting Yb. Cell Rep. 2015;12(3):429–40.

64. Saito K, Inagaki S, Mituyama T, Kawamura Y, Ono Y, Sakota E, et al. A regulatory circuit for piwi by the large Maf gene traffic jam in Drosophila. Nature. 2009;461(7268):1296–9.

65. Pelisson A, Sarot E, Payen-Groschene G, Bucheton A. A novel repeat-associated small interfering RNA-mediated silencing pathway downregulates complementary sense gypsy transcripts in somatic cells of the Drosophila ovary. J Virol. 2007;81(4):1951–60.

66. Yashiro R, Murota Y, Nishida KM, Yamashiro H, Fujii K, Ogai A, et al. Piwi Nuclear Localization and Its Regulatory Mechanism in Drosophila Ovarian Somatic Cells. Cell Rep. 2018;23(12):3647–57.

67. Schwartz LM, Brown C, McLaughlin K, Smith W, Bigelow C. The myonuclear domain is not maintained in skeletal muscle during either atrophy or programmed cell death. Am J Physiol Cell Physiol. 2016;311(4):C607–C15.

68. Lockshin RA, Williams CM. Programmed cell death—I. Cytology of degeneration in the intersegmental muscles of the Pernyi silkmoth. Journal of insect physiology. 1965;11(2):123–33.

69. Finlayson L. Normal and induced degeneration of abdominal muscles during metamorphosis in the Lepidoptera. Journal of Cell Science. 1956;3(38):215–33.

70. Wyllie AH, Kerr JF, Currie AR. Cell death: the significance of apoptosis. Int Rev Cytol. 1980;68:251–306.

71. Schwartz LM, Smith SW, Jones ME, Osborne BA. Do all programmed cell deaths occur via apoptosis? Proc Natl Acad Sci U S A. 1993;90(3):980–4.

72. Berger NA. Poly(ADP-ribose) in the cellular response to DNA damage. Radiat Res. 1985;101(1):4–15.

73. Li W, Jin Y, Prazak L, Hammell M, Dubnau J. Transposable elements in TDP-43-mediated neurodegenerative disorders. PLoS One. 2012;7(9):e44099.

74. Tan H, Qurashi A, Poidevin M, Nelson DL, Li H, Jin P. Retrotransposon activation contributes to fragile X premutation rCGG-mediated neurodegeneration. Hum Mol Genet. 2012;21(1):57–65.

75. Jonsson ME, Garza R, Johansson PA, Jakobsson J. Transposable Elements: A Common Feature of Neurodevelopmental and Neurodegenerative Disorders. Trends Genet. 2020;36(8):610–23.

76. Reilly MT, Faulkner GJ, Dubnau J, Ponomarev I, Gage FH. The role of transposable elements in health and diseases of the central nervous system. J Neurosci. 2013;33(45):17577–86.

77. Dubnau J. The Retrotransposon storm and the dangers of a Collyer’s genome. Curr Opin Genet Dev. 2018;49:95–105.

78. Madigan JP, Chotkowski HL, Glaser RL. DNA double-strand break-induced phosphorylation of Drosophila histone variant H2Av helps prevent radiation-induced apoptosis. Nucleic Acids Res. 2002;30(17):3698–705.

79. Schwartz LM, Truman JW. Hormonal control of muscle atrophy and degeneration in the moth Antheraea polyphemus. J Exp Biol. 1984;111:13–30.

80. Langmead B, Trapnell C, Pop M, Salzberg SL. Ultrafast and memory-efficient alignment of short DNA sequences to the human genome. Genome Biol. 2009;10(3):R25.

81. Kim D, Pertea G, Trapnell C, Pimentel H, Kelley R, Salzberg SL. TopHat2: accurate alignment of transcriptomes in the presence of insertions, deletions and gene fusions. Genome Biol. 2013;14(4):R36.

